# Lipid anchoring and electrostatic interactions target the phospholipase NOT-LIKE-DAD to pollen endo-plasma membrane

**DOI:** 10.1101/2020.10.05.326157

**Authors:** Laurine M. Gilles, Veronica La Padula, Nathanaël M.A. Jacquier, Jean-Pierre Martinant, Peter M. Rogowsky, Thomas Widiez

## Abstract

Phospholipases are ubiquitous enzymes that cleave phospholipids, one major constituent of membranes. They are thus essential for many developmental processes, including male gamete development. In flowering plants, mutation of phospholipase NOT-LIKE-DAD (NLD) leads to peculiar defects in sexual reproduction. Indeed, pollination of a wild-type female with mutant pollen generates haploid embryos containing solely maternal genetic information. Contrary to previous reports NLD does not localize to cytosol and plasma membrane (PM) of sperm cells but to the pollen endo-plasma membrane (endo-PM), a specific membrane derived from the PM of the pollen vegetative cell that encircles the two sperm cells. Pharmacological approaches coupled with targeted mutagenesis revealed that lipid anchoring together with electrostatic interactions between membrane and NLD are involved in the attachment of NLD to this atypical endo-PM. Membrane surface-charge and anionic lipid bio-sensors indicated that phosphatidylinositol-4,5-bisphosphate (PIP(4,5)P_2_) is enriched in the endo-PM as compared to the PM. Our results uncover a unique example of how membrane electrostatic properties can specify a unique polar domain (i.e. endo-PM), which is critical for plant reproduction and gamete formation.

## Introduction

Sexual reproduction represents an evolutionary success and it is thus widespread among eukaryotes (Otto & Lenormand, 2002). It allows mixing genetic information from parents of opposite sexes. Phospholipases are ubiquitous enzymes that hydrolyze phospholipids, a major component of cellular membranes, and are implicated in multiple cellular processes (Park *et al,* 2012; Wang, 2001; Burke & Dennis, 2008). Thus it is not surprising to find evidence that phospholipases are important for sexual reproduction in numerous organisms (Kelliher *et al,* 2017; Gilles *et al,* 2017a; Roldan, 2007; Fry *et al,* 1992; Barman *et al,* 2018). Unlike animals, a double fertilization event is required in flowering plants to achieve sexual reproduction giving rise to the embryo and a nourishing tissue, the endosperm (Walbot & Evans, 2003; Dresselhaus *et al,* 2016). A second distinctive feature from animal systems is that flowering plants do not have motile sperm cells and that the sperm cells are passively carried to the egg apparatus by the pollen tube (Dresselhaus *et al*, 2016). Finally, plant sperm cells are not the direct product of meiosis but stem from two successive mitoses of the initial haploid cell (microspore) (McCormick, 2004; Zhou *et al,* 2017a; Hackenberg & Twell, 2019). The first post-meiotic division (pollen mitosis I) is asymmetric and leads to the formation of the vegetative cell, which does not divide further, and a smaller generative cell. This generative cell goes through an additional mitosis (pollen mitosis II) to produce two sperms cells. Mature pollen (the male gametophyte) is composed of three haploid cells: a large vegetative cell that engulfs two sperm cells (**Fig 1A**). Although small in size, a gametophyte thus exhibits a unique topology having cells within a cell (McCue *et al,* 2011). Yet, how this contributes to fertilization is largely unknown.

**Figure 1.**
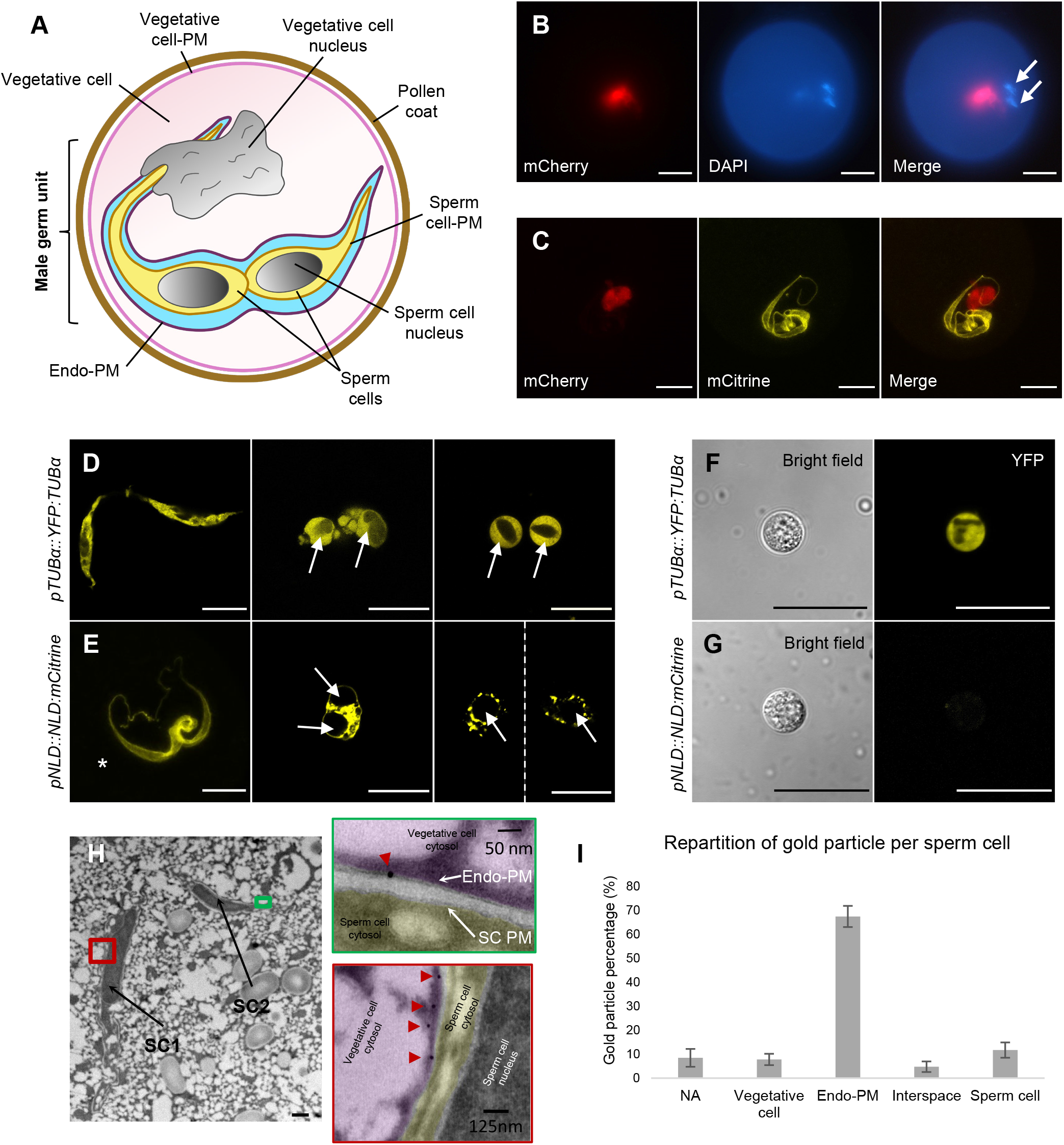
*NLD* is expressed in the vegetative cell of maize pollen and protein localizes to the endo-plasma membrane (endo-PM) of the Male Germ Unit (MGU). **A** Schematic representation of the mature pollen grain organization, focusing on the MGU structure. The two sperm cells (yellow color) are surrounded by an endo-PM (dark purple) originating from the PM of the vegetative cell. **B** Characterization of *NLD* promoter activity (*pNLD::H2B5:mCherry*) in mature pollen. Promoter activity (red signal) is found in the vegetative cell nucleus and not in the sperm cell nuclei. Nuclei are stained with DAPI (4’6-diamidino-2-phenylindole) (blue signal), white arrows indicate the sperm cell nuclei. **C** Subcellular localization of NLD protein (yellow signal) together with *NLD* promoter activity (red signal) in mature pollen. Confocal imaging (Z-stack projection) of mature pollen expressing *pNLD::NLD:mCitrine* and *pNLD::H2B5:mCherry* constructs shows that NLD:mCitrine protein localizes at the MGU, whereas the promoter activity is found in the nucleus of the vegetative cell. **D - G** Following of sperm cells released from mature pollen of pTUBα::YFP:TUBα (D,F) and pNLD::NLD:mCitrine (E,G) lines after osmotic choc (D,E) and sperm cell isolation using Percoll gradient (F,G). YFP:TUB*α* fusion protein is localized inside the sperm cells, while NLD:mCitrine fusion protein is not detected inside isolated sperm cells. E, asterisk* show a MGU before pollen burst. Scale bars: B-G = 20 μm **H, I** Immunogold labeling of NLD:citrine protein in the MGU. (H) Transmission electron microscopy (TEM) pictures of mature pollen of the *pNLD::NLD:mCitrine* line focusing on the ultrastructure of the two sperm cells. The red and green boxes are close up views to visualize gold particles (red arrow heads) that recognize anti-citrine antibody and localize to the endo-PM of the MGU, and not the PM of sperm cell (SC PM). Scale bar in H = 0.5 μm. **I** Quantification of gold particles supporting preferential localization of NLD:mCitrine fusion protein at the endo-PM. Vegetative cell: particles found within the vegetative cell. Endo-PM: particles found in the Endo-PM. Interspace: particles found within the space between sperm cell PM and endo-PM; Sperm cell: particles found in sperm cells (cytoplasm, nucleus and sperm cell-PM). NA: particles that could not be assigned to a particular compartment.

The lack of certain phospholipases impairs pollen development (Kelliher *et al*, 2017; Gilles *et al*, 2017a; Kim *et al*, 2011; Liu *et al*, 2017) and, in the case of a particular maize mutant, leads to the formation of haploid embryos with only the chromosomes from the female parent, lacking the ones from the mutant male parent (Coe, 1959; Barret *et al,* 2008; Jacquier *et al,* 2020). This maize mutant not only represents an interesting resource to study pollen development and plant reproduction but also a powerful plant breeding tool (Jacquier *et al,* 2020; Gilles *et al,* 2017b). Maize breeders have capitalized on this reproductive anomaly, through gynogenesis: after genome doubling of haploid embryos/plantlets, breeders can produce “doubled haploid” plants that are homozygous at all loci in only two generations, compared to the traditional 7 generations of backcrossing or selfing (Jacquier *et al,* 2020; Kalinowska *et al,* 2019). The gene *MATRILINEAL / NOT LIKE DAD / ZmPHOSPHOLIPASE-A* (for simplification designated *NLD* hereafter) was recently found to cause the haploid induction (Kelliher *et al,* 2017; Gilles *et al,* 2017a; Liu *et al,* 2017). Sequence analysis revealed that NLD belongs to the patatin-like phospholipase family (Gilles *et al,* 2017a). The patatin catalytic domain is widely spread in bacteria, yeast, plants and animals, and members of this group are involved in diverse cellular functions (Scherer *et al,* 2010; Baulande & Langlois, 2010; Wilson *et al,* 2006; Kienesberger *et al,* 2009). How NLD functions is not understood yet. NLD was reported to localize at the sperm cell plasma membrane (PM) and in the cytoplasm (Gilles *et al,* 2017a; Kelliher *et al,* 2017). In addition, single sperm cell sequencing documented chromosome fragmentation in sperm cells of the *nld* mutant (Li *et al,* 2017), suggesting a direct or indirect effect of NLD on genome stability. The original *nld* mutant carries a 4-bp insertion in the last exon of *NLD*. This mutation causes a frameshift that replaces the last 49 amino acids of the wild-type protein by 20 unrelated amino acids followed by a premature STOP codon (Gilles *et al,* 2017a; Kelliher *et al,* 2017). The resulting truncated protein is unstable and does not accumulate in maize pollen, whereas in the heterologous *Arabidopsis thaliana* (designated *Arabidopsis* hereafter) root cell system it loses PM localization (Gilles *et al*, 2017a; Kelliher *et al*, 2017). These results indicate a critical role of the carboxy-terminus (C-ter) of NLD, but the mechanism through which the C-ter truncation of NLD leads to loss of protein stability and haploid induction is not known.

Here we demonstrate NLD does not localize to sperm cell PM but, instead, to the endo-PM that surrounds the sperm cells. This endo-PM belongs to the adjacent vegetative cell. We demonstrate that the C-ter part of the NLD protein is not sufficient by itself to ensure subcellular localization, but it comprises important amino acid residues, which, together with additional amino acid residues in the N-ter, are key to target NLD to the PM. Pharmacological and mutagenesis approaches demonstrated the involvement of both lipid anchoring and electrostatic interactions to properly address NLD to the pollen endo-PM.

## Results and Discussion

### *NLD* promoter is active in the vegetative cell of maize pollen

In mature maize pollen, the vegetative cell fully encloses the two sperm cells forming a complex cellular structure named the male germ unit (MGU), which regroups all the genetic material of the pollen grain (**Fig 1A**) (McCue *et al,* 2011; Dumas *et al,* 1984). The sperm cells are thus “cells within a cell”. Previous studies reported the sub-cellular localization of the NLD protein, expressed under its own promoter, in both plasma membrane (PM) and cytoplasm of sperm cells (Gilles *et al*, 2017a; Kelliher *et al,* 2017). In order to refine the *NLD* spatial and temporal expression pattern during maize pollen development, we developed a new promoter reporter line *pNLD::H2B:2x-mCherry.* It contains the same 2.6-kbp *NLD* promoter fragment used in our previous study (Gilles *et al,* 2017a) driving the expression of two mCherry fluorescent proteins that are fused to histone H2B in order to target fluorescence to nuclei. Imaging of mature pollen grains surprisingly revealed promoter activity in the vegetative cell nucleus, and not the sperm cell nuclei as one would have expected based on previous observations of NLD protein localization (**Fig 1B**). This observation was confirmed using eight independent transgenic lines (**Appendix Table S1**). It is further supported by *NLD* expression data in a recent transcriptomic data set from isolated maize sperm cells (Chen *et al,* 2017), which indicates the absence of *NLD* transcripts from sperm cells. Time course experiments during microgametogenesis demonstrated that the *NLD* promoter activity starts during the late bicellular stage, after the DNA condensation of the generative cell, just prior to the second pollen mitosis, which leads to the formation of two sperm cells after mitotic division of the generative cell (**Fig EV1**). The start of *NLD* expression at late bicellular stage is consistent with previous observations based on both GUS promoter activity (but lacking subcellular resolution) (Gilles *et al,* 2017a), and RNA-seq kinetics on whole pollen grain (Liu *et al,* 2017).

### NLD protein localizes specifically to the endo-plasma membrane of the vegetative cell that surrounds the sperm cells

Pollen grains expressing both the *pNLD::H2B:2x-mCherry* promoter fusion and the *pNLD::NLD:mCitrine* protein fusion underline a conflicting situation in which the *NLD* promoter is active in the vegetative cell (red color in **Fig 1C**), whereas the NLD protein seems to be localized in the adjacent sperm cells (yellow color in **Fig 1C**). Complementation assays demonstrated that the NLD:mCitrine protein fusion is functional since it fully abolished the haploid induction phenotype of the *nld* mutant (**Appendix Table S2**). Two possible explanations of this uncommon pattern were that (1) *NLD* transcripts (or proteins) are synthesized in the vegetative cell and then move to the sperm cells as reported for *ABA-hypersensitive germination 3 (AHG3)* transcript in *Arabidopsis* (Jiang *et al,* 2015), or (2) NLD protein is not localized within the sperm cells, but rather at the endo-plasma membrane (endo-PM) of the vegetative cell, which tightly surrounds the sperm cells (**Fig 1A**) (Li *et al,* 2013; McCue *et al,* 2011; Dumas *et al,* 1984).

To test these hypotheses, we simplified the experimental system by disconnecting the sperm cells from the male germ unit including the endo-PM. Both the *pNLD::NLD:mCitrine* line and a control sperm cell marker line, for which α-tubulin-YFP (pTUBα::YFP:TUBα) had been demonstrated to be expressed in sperm cells and not in the vegetative cell (Kliwer & Dresselhaus, 2010), were submitted to osmotic shock directly under a microscope slide and imaged during the process of sperm cell release (**Fig 1D, E**). The α-tubulin-YFP line marked evenly the sperm cell cytosol and allowed to clearly visualize the isolated sperm cell (**Fig 1D**). In contrast, the application of hyperosmotic shock to *pNLD::NLD:mCitrine* pollen gave rise to a citrine fluorescent signal, which first surrounded the sperm cells and was then progressively lost when sperm cells individualized (**Fig 1E**). These observations were confirmed by a complementary approach of sperm cell isolation using a Percoll gradient (Dupuis *et al,* 1987) (**Fig 1F, G**). Sperm cells derived from the α-tubulin-YFP reporter line allowed to clearly distinguish the individualized sperm cells based on YFP fluorescence signal, whereas no fluorescent was detected in sperm cells derived from *pNLD::NLD:mCitrine* pollen. These results are in favor of the hypothesis that NLD localizes to the endo-PM of the vegetative cell. In mature pollen, the close proximity, the twisted nature of the MGU and the strength of the signal likely gave the erroneous impression of a localization within the sperm cells.

To fully confirm this hypothesis, immuno-electron microscopy (immuno-EM) experiments were performed on ultrathin sections (100 nm) of pollen grains containing functional *pNLD::NLD:mCitrine* construct. NLD was labeled using an antibody against mCitrine followed by antibody detection with a Protein A-gold electron dense probe. Non-transgenic mature pollen grains followed the same protocol as an internal control to assess background level in the transmission electron microscope (TEM). Observation of the immuno-EM staining of transgenic *(pNLD::NLD:mcitrine)* and control pollen grains revealed the ultrastructure of MGU and an enrichment of gold particles in the endo-PM surrounding the sperm cells and not in the PM of the sperm cells **(Fig 1H**). Quantification of different pollen grains demonstrated that the NLD:mCitrine fusion protein localizes predominantly at the endo-PM, clarifying NLD subcellular localization (**Fig 1I, Appendix Table S3**). This vegetative endo-PM is extremely poorly characterized and even often omitted from text books. To our knowledge only the small GTP-binding Rho of Plants (ROP9) protein has been reported to have endo-PM localization in *Arabidopsis* (Li *et al,* 2013, 9; Lin *et al,* 1996). In terms of plant breeding, considering that haploid induction through NLD is important for maize breeding (Jacquier *et al,* 2020), physical, chemical or other treatments to interfere with the properties of the endo-PM could be envisioned as a means for haploid induction.

### Endo-PM NLD localization is conserved in *Arabidopsis* pollen

To further investigate NLD association to the endo-PM, the *Arabidopsis* plant model was used as a heterologous system to save time, since the development of transgenic maize lines takes one year and the seed to seed life cycle is more than twice longer than in *Arabidopsis.* To drive gene expression in the vegetative cell and not in the sperm cells, similarly to the maize *NLD* promoter in pollen, the promoter of the *POLYUBIQUITIN 10* gene *(At4G05320)* was chosen. Although *AtUBQ10* is transcribed in nearly all plant tissues, it has been reported to show high expression in pollen grain (thus containing both vegetative and sperm cells), but low expression in isolated sperm cell (Borges *et al,* 2008). In addition, the high promoter activity in roots allowed the use of the same *Arabidopsis* lines to carry out confocal observations in a more conventional plant cell system without endo-PM. The expected *AtUBQ10* promoter pattern was experimentally validated by confocal imaging of a *pUBQ10::VENUS-N7* transgenic line targeting the VENUS fluorescent protein to the nucleus (**Fig EV2**). In the pollen context, an expression pattern specific of the vegetative cell was demonstrated and no fluorescence was detected in sperm cell (**Fig EV2A**), similarly to what had been found for the *NLD* promoter in maize pollen (**Fig 1B**). In addition, the promoter was indeed found to be active in root cells (**Fig EV2B**), as reported earlier (Geldner *et al,* 2009; Grefen *et al,* 2010).

Previous work in the heterologous *Arabidopsis* root expression system had shown that NLD full-length protein (NLD_full_, **Fig 2A**) localizes predominantly to the PM, but also to intracellular compartment and cytosol (Gilles *et al,* 2017a), whereas the truncated NLD protein (NLD_trunc_; **Fig 2A**), which is responsible for haploid induction phenotype, is not recruited at the PM (Gilles *et al,* 2017a). We confirmed these previous results in root tip cells (**Fig 2E, F**) and quantified by measuring the ratio of fluorescence found at the PM versus fluorescence found in the cytosol (**Fig 1L**). Whereas NLD_full_:mCitrine had a PM/cyto fluorescence ratio of 2.71 indicating PM enrichment over cytosol, NLD_trunc_:mCitrine had a PM/cyto fluorescence ratio of 0.71. In *Arabidopsis* pollen grains, NLD_full_:mCitrine displays a strong fluorescent signal at the MGU and was not detected at the PM (**Fig 2B**). This signal likely corresponds to the endo-PM as for maize. The truncation of NLD (NLD_trunc_:mCitrine) provoked loss of this subcellular pattern, with fluorescent signal found only in the cytosol and in intracellular compartments of the vegetative cell (**Fig 2C**). Of note, the weaker fluorescent signal surrounding the pollen grain was likely due to auto-fluorescence from the pollen coat since similar auto-fluorescence was observed in pollen grain of the non-transgenic Col-0 control line (**Fig 2D**). Taken together these observations suggest that NLD protein subcellular localization is highly conserved between maize and *Arabidopsis* pollen and that NLD truncation leads to a loss membrane localization both to the PM in roots and the endo-PM in pollen.

**Figure 2.**
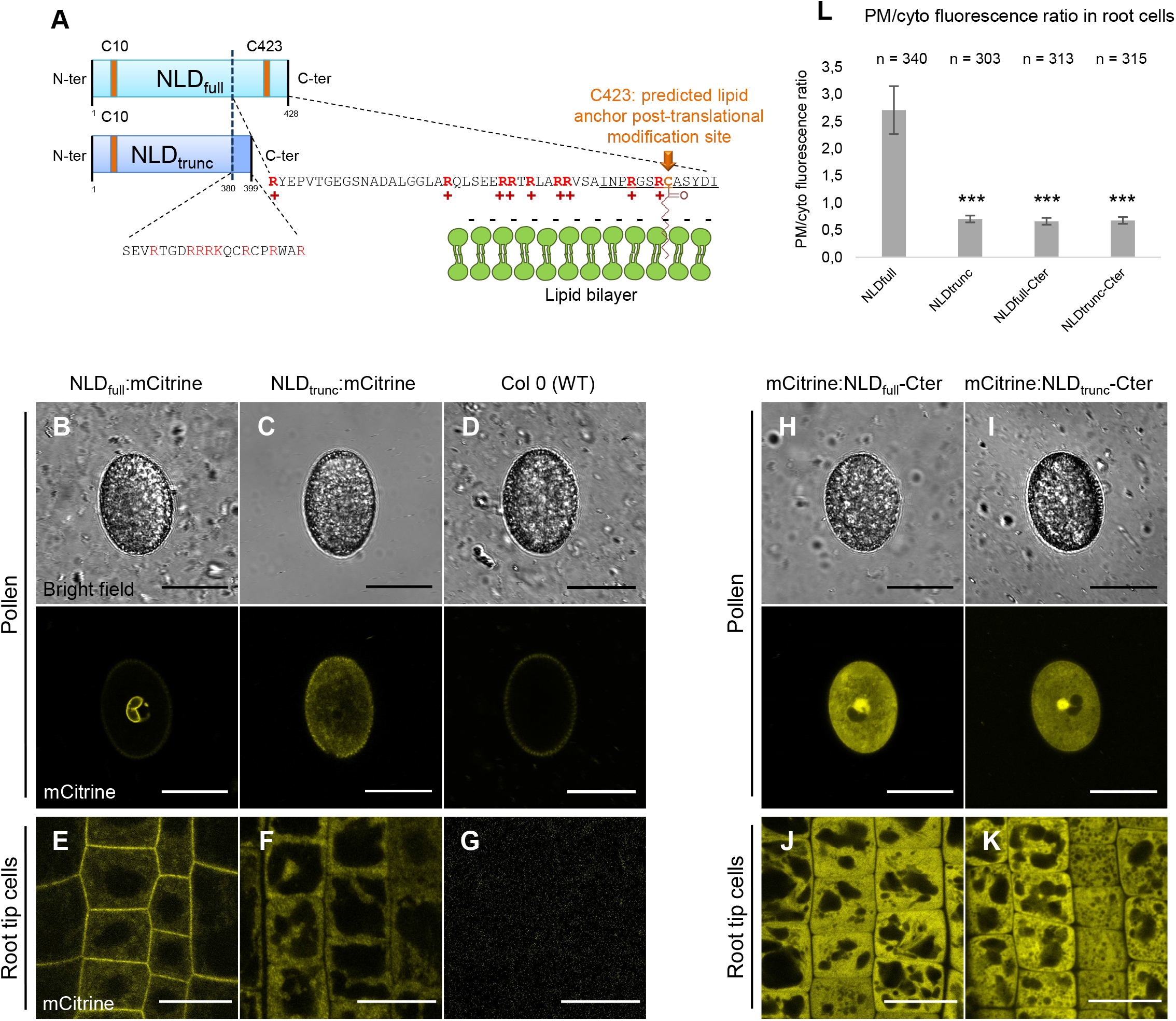
NLD protein localization is conserved in *Arabidopsis* pollen. **A.** Schematic representation of NLD full-length (NLD_full_) and truncated (NLD_trunc_) proteins. Since the NLD C-ter region contains both a predicted lipid anchor post-translational modification site (C423) and a stretch of positively charged amino acids (in red in the sequence), targeting of NLD to membrane could be achieved *via* lipid anchoring and/or *via* electrostatic interactions between the anionic lipids present at the membrane surface and the cationic amino acid present at the NLD C-ter. **B-K.** Confocal imaging of different forms of NLD protein in *Arabidopsis* pollen and root cells. Confocal imaging of mature pollen (B, C, D, H, I) and root tips (E, F, J, K) from non-transformed Columbia 0 (Col 0) (D, G) and lines expressing the full length NLD protein (NLD_full_:mCitrine) (B,E), the truncated NLD protein (NLD_trunc_:mCitrine) (C, F), the C-terminal parts of NLD_full_ (mCitrine:NLD_full_-Cter) (H, J), and the C-terminal parts of NLD_trunc_(mCitrine:NLD_trunc_-Cter) (I, K). Scale bars: root cells = 10 μm, mature pollen = 20 μm. **L**. Quantification of mutagenesis effect on localization using the ratio of the fluorescence intensity at plasma membrane compared to the florescence intensity in cytosol (PM/cyto fluorescence ratio) in root cells (mean ± SD). *** denotes significant difference between means as compared to NLD_full_:mCitrine (p-value < 0.001; Wilcoxon test, n = cells number analyzed).

### The C-terminal part of the NLD protein is not sufficient to ensure subcellular localization

The reversible addition of a fatty acid with 16 carbon atoms (palmitate) to a protein is called S-palmitoylation or, more generally, S-acylation. This post-translational modification is emerging as a ubiquitous mechanism to control membrane affinity, trafficking, maturation and function of proteins (Chamberlain & Shipston, 2015; Hurst & Hemsley, 2015). Two cysteine residues (C10 and C423) of the NLD protein sequence were predicted *in silico* to be S-acylated (**Fig 2A**) (Gilles *et al,* 2017a). In addition, a stretch of positively charged amino acid residues (Arginine, R), was found in close proximity of the predicted C423 lipid anchor site (**Fig 2A**). These *in silico* observations fed the working hypothesis that NLD was specifically addressed to the endo-PM *via* a lipid anchor and/or electrostatic interactions between positively charged amino acids of NLD and negatively charged membrane lipids (**Fig 2A**). Similar protein/membrane interactions have been demonstrated to operate in other cellular context in plant (Hurst & Hemsley, 2015; Alassimone *et al,* 2016; Simon *et al,* 2016; Platre *et al,* 2018; Barbosa *et al,* 2016), or animal cells (Chamberlain & Shipston, 2015; Zhou *et al,* 2017b; Liu *et al,* 2007; He *et al,* 2007). Since NLD_trunc_ loses membrane localization (PM in root cells and endo-PM in pollen) and since all amino acids concerned by this hypothesis are located within the C-ter part, except the C10, we investigated whether the C-ter of NLD_full_ is sufficient to ensure membrane localization. The NLD sequence corresponding to the last 49 amino acids that are absent in NLD_trunc_, was fused to mCitrine (mCitrine:NLD_full_-Cter), just like the unrelated sequence of 20 amino acids replacing it in NLD_trunc_ (mCitrine:NLD_trunc_-Cter). Confocal imaging of pollen grain of both lines showed fluorescent signal in the cytosol and nucleus of the vegetative cell, and absence of fluorescent in sperm cells (**Fig 2H and I**). Cytosolic fluorescence was also observed in root cells (**Fig 2J and K**), and PM/cyto fluorescence ratios (0.66 and 0.68, respectively) substantiated quantitatively cytosolic enrichment as compared to PM (**Fig 2H**). These observations suggest that the C-ter part of NLD_full_ is not sufficient to address the mCitrine fluorescent protein to membranes, indicating that other region(s) of NLD contribute to its PM localization. The presence of 8 positively charged amino acids in the unrelated C-ter of NLD_trun_c further underlined that other features than solely positive charged amino acids are required for the proper NLD localization.

### Membrane attachment of NLD protein is mediated by S-acylation

Multiple reports, in both animals and plants, show that a lipid anchor allows membrane attachment for proteins that do not contain transmembrane domains (Chamberlain & Shipston, 2015; Hurst & Hemsley, 2015). Thus, the involvement of the two predicted lipid anchor sites C10 and C423 (**Fig 2A**) was first examined by a pharmacological approach. 2-bromopalmitate (2-BP), an inhibitor of S-Acyl transferase which catalyzes the S-acylation reaction (Webb *et al*, 2000), was used to treat 7-day-old *Arabidopsis* seedlings expressing NLD_full_:mCitrine (**Fig 3**). Confocal imaging of root cells after a 2 h treatment with 50 μM 2-BP revealed a partial NLD_full_:mCitrine delocalization from the plasma membrane to the cytosol (**Fig 3B**) as compared to mock treatment (**Fig 3A**). Fluorescent intensity quantification confirmed this observation, the PM/cyto fluorescence ratio significantly decreasing for root cells after 2-BP treatment as compared to mock treatment (2.79 to 0.99) (**Fig 3E**). To ensure that the 2-BP treatment had no major effects on PM integrity, similar 2-BP treatment was applied to seedlings expressing the PM localized GFP:LTI6b construct, where the fusion protein is inserted in the PM by two transmembrane domains (Platre *et al,* 2018; Cutler *et al,* 2000) (**Fig 3C and D**). The 2-BP treatment had no effect on GFP:LTI6b subcellular localization, the signal being maintained at the PM (**Fig 3C-F**). Pollen expressing NLD_full_:mCitrine in the vegetative cell was also treated with 2-BP. Similar treatment as done for root cells (50 μM for 2h) did not affect NLD localization in pollen, most certainly because the protective pollen coat, which forms an impermeable barrier, partially impedes drug penetration. Increasing the 2-BP concentration to 150 μM resulted in significantly less pollen grains having NLD_full_:mCitrine localized to the endo-PM (13.4% in 2-BP treated vs. 80.8% in mock treated pollen) and more pollen grains were found to have either spotty or cytosolic NLD_full_:mCitrine signal as compared to the pollen treated with mock (**Fig EV3**). Altogether, these results demonstrate the involvement (direct or indirect) of S-acylation in the stable attachment of NLD to the PM in root cells and to the endo-PM in pollen.

**Figure 3.**
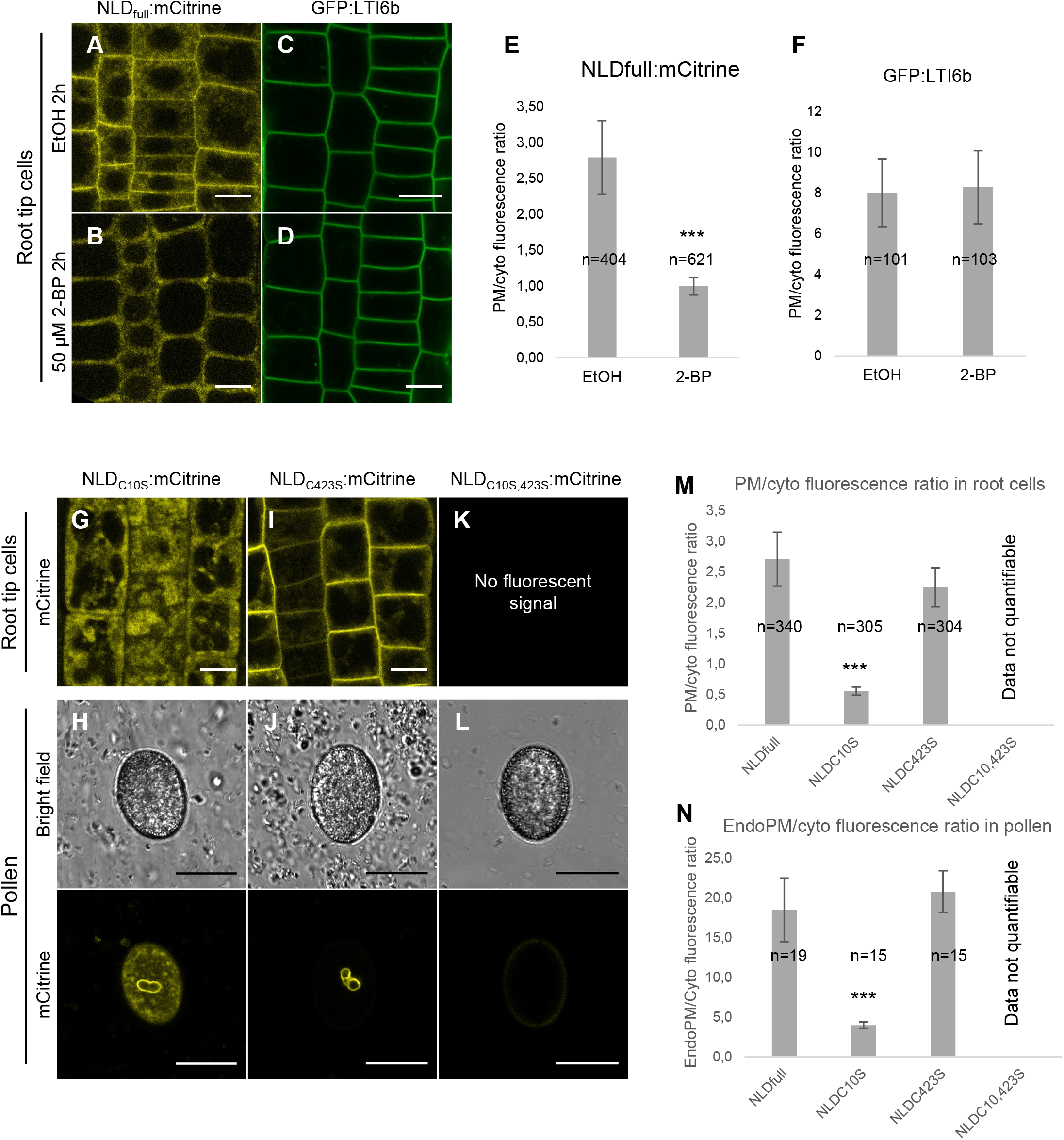
Lipid anchors are necessary for NLD attachment and stability at membranes. **A - F.** Pharmacological approach showing the involvement of S-palmitoylation in NLD addressing to the PM of root cells. Confocal imaging of root cells expressing NLD_full_:mCitrine (A, B) or transmembrane GFP:LTI6B control (C,D) after 2 h treatment with 50 μM 2-BP (2-bromopalmitate), an inhibitor of palmitoylation (B, D), or with mock treatment (EtOH) (A, C). (E, F) Quantification of drug treatment effect on NLD subcellular localization using the ratio of the fluorescence intensity at plasma membrane compared to the fluorescence intensity in cytosol (PM/cyto fluorescence ratio) in root cells (mean ± SD). *** denotes significant difference (p-value < 0.001; Wilcoxon test, n = cells number analyzed). Scale bars: root cells = 10 μm. **G - L.** Targeted mutagenesis of predicted NLD lipid anchor sites C10 and C423. Confocal imaging of root cells and mature pollen from lines: NLD_c10S_:mCitrine (C10 mutated to serine (S)) (G,H), NLD_C423S_:mCitrine (I,J) and NLD_C10S423S_:mCitrine (K,L). Scale bars: pollen grains = 20 μm. **M, N** Quantification of mutagenesis effect on localization using the ratio of the fluorescence intensity at plasma membrane compared to the florescence intensity in cytosol (PM/cyto fluorescence ratio) in root cells (M) and in pollen grain (N) (mean ± SD). *** denotes significant difference (p-value < 0.001; Wilcoxon test, n = cells number analyzed).

To further support NLD membrane attachment by S-acylation of cysteine residues, targeted mutagenesis was carried out on C10 and C423 (**Fig 2A**). The C residues were separately or simultaneously substituted by serine (S) residues leading to the following constructs: NLD_C10S_:mCitrine, NLD_C423S_:mCitrine and NLD_C10S,C423_s:mCitrine (**Fig 3G-N**). Confocal imaging of root cells revealed drastic delocalization of NLD_C10S_:mCitrine from the PM to the cytosol and intracellular compartments, with a significant decrease of the PM/cyto fluorescent ratio from 2.71 for NLDfull:mCitrine to 0.56 for NLDC10S:mCitrine (**Fig 3G and M**). The NLDC423S:mCitrine fusion protein remained accumulated at the PM with no significant change in the PM/cyto fluorescent ratio as compared to NLD_full_:mCitrine (2.25 vs 2.71, respectively) (**Fig 3I and M**). The simultaneous mutation of both C10 and C423 resulted in a loss of fluorescent signal (**Fig 3K and M**) in 15 independent lines (**Appendix Table S1**). These results underline the importance of these two cysteine residues in promoting NLD attachment to membrane, which are likely important to stabilize the protein based on evidences gained in other biological contexts (Chamberlain & Shipston, 2015; Hurst & Hemsley, 2015). Similar conclusions on the importance of both C10 and C423 for correct NLD localization and stability could be drawn based on observations of these three mutated versions in pollen (**Fig 3H, J, L and N**). Contrary to the root results, NLD_C10S_:mCitrine showed only partial delocalization of fluorescent signal from the endo-PM (**Fig 3H and N)**, whereas it was almost fully removed from PM in root cells (**Fig 3G and M**), indicating that NLD is probably more strongly attached to endo-PM as compared to root cell PM. Altogether, both pharmacological and targeted mutagenesis approaches demonstrated the importance of a lipid anchoring mechanism to stabilize NLD at membranes, at the PM in roots and specifically at the endo-PM in pollen. Interestingly, for the only other known endo-PM localized protein ROP9, three cysteine residues have been shown to be critical for proper endo-PM localization (Li *et al,* 2013, 9).

### NLD targeting to membranes involves electrostatic interactions

If S-acylation is often essential for anchoring proteins within lipid bilayers (membranes), the specific attachment to particular membranes can be fine-tuned through electrostatic interactions with lipids (Simon *et al,* 2016; Platre *et al,* 2018; Zhou *et al,* 2017b). To address the importance of membrane electrostatics for the subcellular localization of NLD, the electronegativity of the membrane surface in cells was indirectly decreased by phenylarsin oxide (PAO) treatment. PAO is an inhibitor of PI4-kinases (PI4Ks), involved in the biosynthesis of highly electronegative PI4P phospholipids, which are predominantly present on the PM surface in root cells (Simon *et al,* 2016, 2014). Confocal imaging of root cells from NLD_full_:mCitrine seedlings after 30 min of PAO treatment at 60 μM showed a clear delocalization of the signal from PM to cytosol and endomembrane compartments (**Fig 4A and B)**, although low fluorescent signal was maintained at PM. Quantification of PM/cyto fluorescence corroborated a statistically significant reduction of this ratio in PAO treated cells (0.98), as compared to mock treatment (2.39) (**Fig 4I**). The effectiveness of short-term PAO treatment was established for control lines expressing either a “PI4P biosensor” (mCitrine:1x-PH^FAPPI^) or a surface charge biosensor (mCitrine:FARN6+) (Simon *et al,* 2016, 2014). After PAO treatment a strong PI4P biosensor delocalization and a partial charge biosensor delocalization were observed in root cells (**Fig 4D and F**) as compared to mock treatment (**Fig 4C and E**). In addition, the absence of a negative effect of short-term PAO treatment on PM integrity was established using *GFP:LTI6b* seedlings (**Fig 4G, H and J**). In pollen, PAO treatment also led to the delocalization of NLD from the endo-PM, but necessitated longer incubation time (1h) and higher concentration (180 μM), again certainly due to the protective effect of pollen coat. Significantly less pollen grains had NLD_full_:mCitrine localized to the endo-PM (12.4% in PAO treated vs 80.4% in mock treated pollen) and more pollen grains were found to have either spotty or cytosolic NLD_full_:mCitrine signal as compared to the pollen treated with mock (**Fig EV4**). Altogether, this pharmacological approach confirmed the importance of membrane electronegativity for correct NLD targeting.

**Figure 4.**
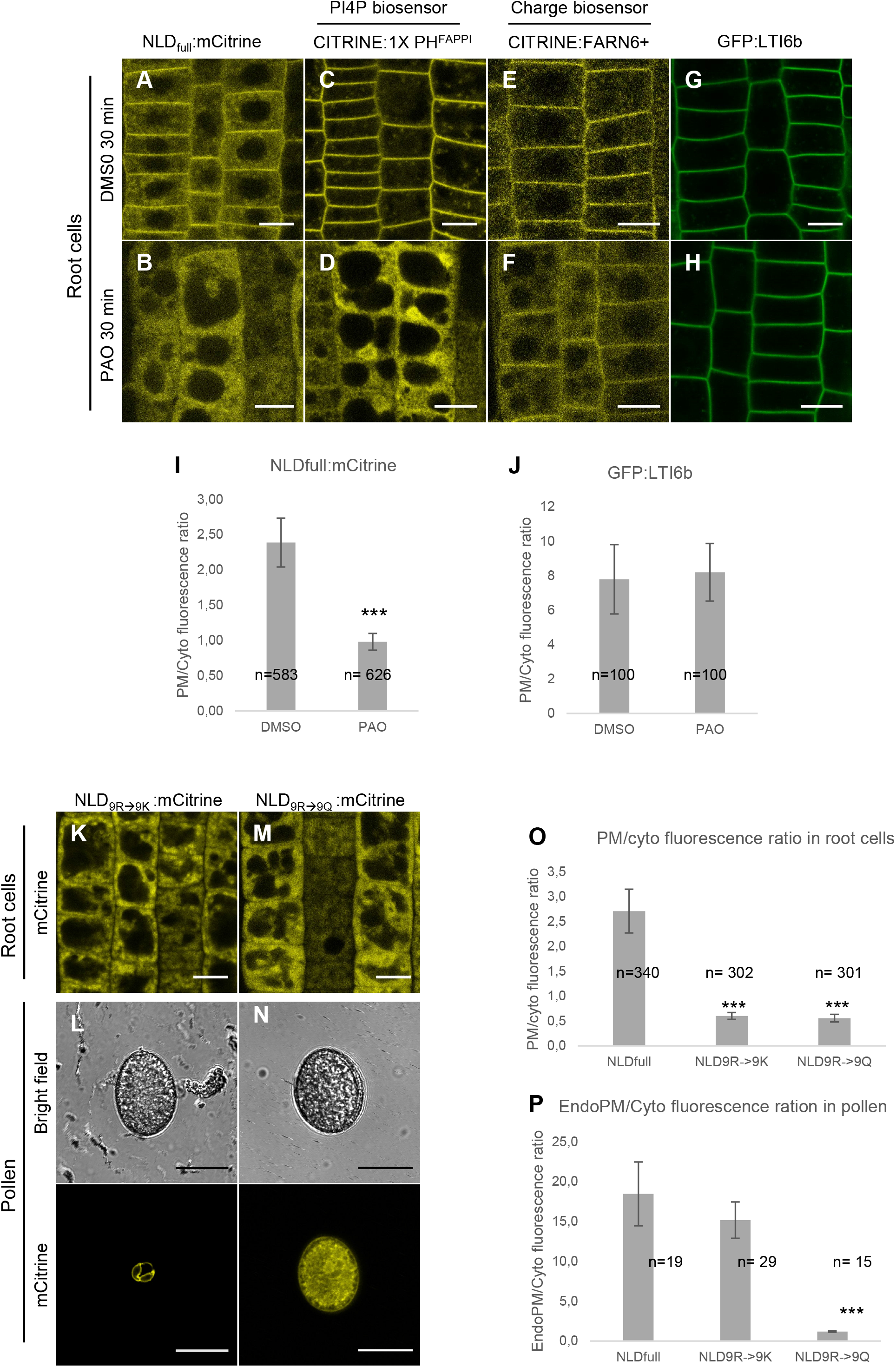
Electrostatic interactions between NLD and negatively charged membranes are necessary for correct targeting. **A-H** Pharmacological approach showing the importance of membrane electronegativity in targeting NLD to PM of root cells. Confocal imaging of root cells from lines expressing NLD_full_:mCitrine (A, B), the PI4P biosensor mCitrine:1XPH^FAPPI^ (C,D), 6+ the biosensor of surface membrane charge mCitrine:FARN (E,F) or transmembrane GFP:LTI6b (G,H) after 30 min treatment with 60 μM PAO (Phenylarsine oxide), an inhibitor of PI4-Kinases (A, C, E, G) or with mock (DMSO) (B, D, F, H). Scale bars: root cells = 10 μm. **I, J**: Quantification of drug treatment effect on NLD subcellular localization using the ratio of the fluorescence intensity at PM vs the fluorescence intensity in cytosol (Fluo PM/cyto) in root cells (mean ± SD). *** denotes significant difference (p-value < 0.001; Wilcoxon test, n = cells number analyzed). **K - P** Targeted mutagenesis of the nine positively charged amino acids located in the NLD C-ter impaired NLD subcellular localization. Confocal imaging of roots cells and mature pollen from lines NLD_9R→9Q_:Citrine (9 arginine (R) residues mutated to lysine (K) residues) (K, L), or NLD_9R→9Q_:Citrine (9 R residues mutated to glutamine (Q) residues) (M, N). Scale bars: pollen grains = 20 μm. O, P: Quantification of mutagenesis effect on localization using the ratio of the fluorescence intensity at plasma membrane compared to the florescence intensity in cytosol (PM/cyto fluorescence ratio) in root cells (O) and in pollen grain (P) (mean ± SD). *** denotes significant difference (p-value < 0.001; Wilcoxon test, n = cells number analyzed).

Mechanistically, positively charged parts of NLD could interact with negative membrane charges. Since the loss of NLDtrunc membrane localization was correlated with the loss of a stretch of positively charged amino acid residues in the missing NLD C-ter part (**Fig 2C**), a targeted mutagenesis was performed to assess the role of this stretch in addressing NLD to membranes. In the resulting NLD_9R→9Q_:mCitrine line the 9 positively charged arginine (R) residues were substituted, in NLD_full_, by 9 neutral glutamine (Q) residues. Confocal imaging of NLD_9R→9Q_:mCitrine root cells showed mainly cytosolic fluorescence (**Fig 4M**), with a significant delocalization from PM to cytosol as compared to non-mutated NLD_full_:mCitrine (PM/cyto fluorescence of 0.56 vs 2.71, respectively) (**Fig 4O**). In pollen, a similar effect was observed with a drastic loss of endo-PM localization accompanied by a significant increase of fluorescent signal in cytosol for NLD_9R→9Q_:mCitrine as compared to NLD_full_:mCitrine (**Fig 4N and P**). In a control experiment the 9 R residues were replaced by 9 lysine (K) residues. This conservative change maintaining the positive charges gave rise to the NLD_9R→9K_:mCitrine line. In pollen, NLD_9R→9K_:mCitrine still localized to the endo-PM as NLD_full_:mCitrine, indicating that maintaining the same number of positively charged amino acid on NLD C-ter was sufficient to maintain NLD at the endo-PM (**Fig 4L and P**). Unexpectedly, NLD_9R→9K_:mCitrine was not targeted to the PM in root cells and rather localized in cytosol (**Fig 4K and O**). Altogether, both the pharmacological and the targeted mutagenesis approach demonstrated the implication of electrostatic interactions between NLD and negatively charged membranes to correctly target NLD, especially to pollen endo-PM.

### The endo-PM of the pollen vegetative cell is enriched for PI(4,5)P_2_ lipid sensor

If the above results on lipid anchoring and electrostatic interactions provided insights in the type of interactions between NLD and membrane, they did not answer the question why NLD was targeted specifically to the endo-PM and not to both endo-PM and vegetative cell PM. To investigate the lipid composition of the poorly characterized endo-PM, we took advantage of a charge sensor and a lipid biosensor collection that had been shown to recognize phosphatidic acid (PA), phosphatidylserine (PS), phosphatidylinositol 3-phosphate (PI3P), Phosphatidylinositol 4,5-bisphosphate (PI(4,5)P2) or phosphatidylinositol 4-phosphate (PI4P) lipids in *Arabidopsis* root cells and other cellular contexts (Simon *et al,* 2014, 2016; Platre *et al,* 2018). All sensors presented the advantage to be expressed under the *pUBQ10* promoter that confers expression in the vegetative cell but not the sperm cells in the pollen grain (**Fig 2A, B**). Four of the 6 tested biosensors (PA, PS, PI4P, charge) localized to both PM and endo-PM (**Fig 5A, B, E and F**). The PIP3 biosensor localized to intracellular compartments of the vegetative cell (**Fig 5C**). This pattern is similar to the one reported for the root cell type, where these intracellular compartments have been identified as late endosomes (Simon *et al,* 2014, 2016; Platre *et al,* 2018). Interestingly, the PI(4,5)P_2_ sensor showed strong enrichment at the endo-PM as compared to the PM (**Fig 5D**). This enrichment of PI(4,5)P2 at the endo-PM, which may also reflect a depletion at the PM, is an appealing candidate to represent a signature that allows distinction between the PM and the endo-PM in the vegetative cell context. Since the PI(4,5)P_2_ sensor was shown to be enriched in the root cell PM (Simon *et al,* 2014), both NLD:mCitrine and PI(4,5)P_2_ sensor show very similar localization in these two cell types analyzed. These observations may explain why NLD is found at the PM of root cells, whereas it localizes at the endo-PM in the pollen vegetative cell.

**Figure 5.**
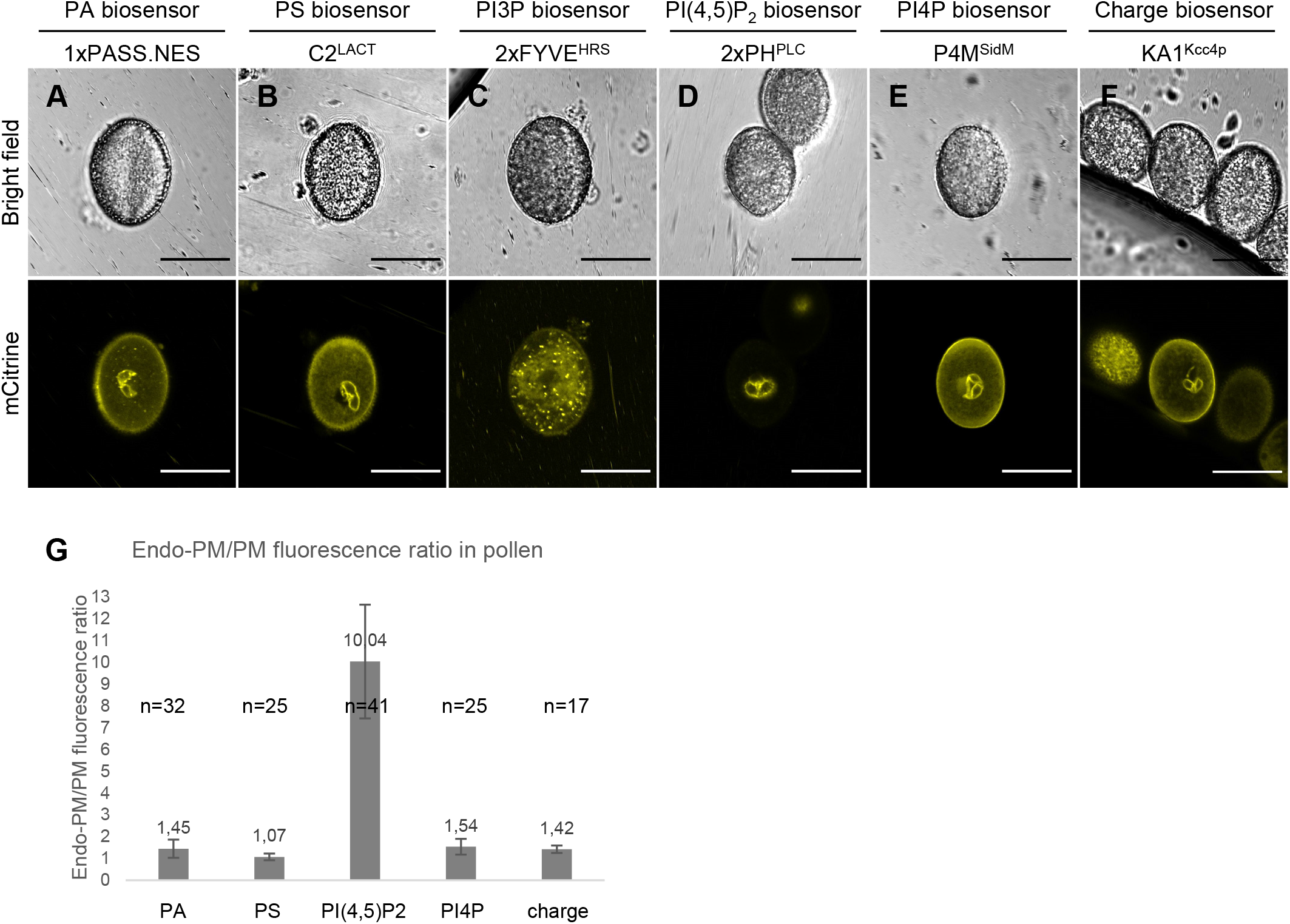
Characterization of endo-PM properties using genetically encoded fluorescent lipid sensors. **A-E** Localization of different lipids in the vegetative cell of *Arabidopsis* mature pollen through lipid biosensors. Confocal imaging allowed to visualize different lipid biosensors (Simon *et al,* 2014; Platre *et al,* 2018, 2019) expressed in the vegetative cell of pollen thanks to pUBQ10 promoter.: PA (phosphatidic acid), PS (phosphatidylserine), PI3P (Phosphatidylinositol-3-phosphate), Pi(4,5)P_2_ (Phosphatidylinositol-4,5-bisphosphate), PI4P (Phosphatidylinositol-4-phosphate). **F** Confocal imaging of *Arabidopsis* mature pollen showing subcellular localization of an anionic amino acid biosensor Kcc4p (KA1^Kcc4p^) which consists in a folded domain that lacks stereo-specificity and associates non-specifically with anionic lipids (Simon *et al*, 2016). **G** Quantification of fluorescence intensity for biosensors at the endo-PM as compared to the fluorescence intensity in PM of vegetative cell (endo-PM/PM fluorescence ration) (mean ± SD). n = number of mature pollens analyzed.

During infection of plant cells by biotrophic pathogens, a feeding structure, called a haustorium, forms inside the plant host cell. Consequently, an extrahaustorial membrane (EHM) derived from the host PM surrounds the haustorium (Hückelhoven & Panstruga, 2011; Gil & Gay, 1977; Roberts *et al*, 1993). Thus, a parallel can be drawn between this EHM in infected cells and the endo-PM in pollen, because both enclose structures of external origin within the cell. The EHM was found to differ in its molecular composition from the conventional PM (Hückelhoven & Panstruga, 2011; Gil & Gay, 1977; Roberts *et al,* 1993; Micali *et al,* 2011; Koh *et al,* 2005). Remarkably, two recent studies reported that the EHM is enriched in PI(4,5)P2 similarly as described in this study for the pollen endo-PM (Shimada *et al,* 2019; Qin *et al,* 2020). Future work on the pollen endo-PM needs to track the genesis of this special interface during male gamete differentiation and to investigate its functional significance. More broadly speaking, the cellular insights of this study lead to the challenge to explain mechanistically the astonishing fact that the loss of NLD outside the sperm cell (on endo-PM) leads to a drastic phenotype on chromosome stability within sperm cells (Li *et al,* 2017). The resulting necessity of interactions between the vegetative pollen cell and the enclosed sperm cells to have proper gamete formation reveals a specific and unique cell-cell communication route, and its dependence on local membrane properties.

## Materials and Methods

### Plant material and plant growth conditions

The maize (Zea mays) plants were cultivated in an S2 greenhouse or S2 growth chambers as previously described (Rousseau *et al,* 2015; Doll *et al,* 2020, 2019). Non-transgenic plants were also cultivated in a field located in Lyon (France). Haploid induction rate was evaluated using female tester line as described elsewhere (Gilles *et al,* 2017a). *Arabidopsis thaliana* plants were grown in soil under long-day conditions at 21°C and 70% humidity. The *in vitro* experiments were done on Murashige and Skoog (MS) Basal Medium supplemented with 0.8% plant agar (pH 5.7) in continuous light conditions at 21°C for 6–9 days. All lines generated in this study and used are presented in **Appendix Table S1**. For generation and growth of all transgenic plants, we followed the national guidelines and legislations.

### Plant transformation and selection

Maize transformation was performed in inbred line A188 (Gerdes & Tracy, 1993). *Agrobacterium*-mediated transformation was executed according to a published protocol (Ishida *et al,* 1996) and T-DNA integrity was checked as described elsewhere (Gilles *et al,* 2017a). *Arabidopsis thaliana* ecotype Col0 was transformed with the floral-dip method which consists to incubate the flowers in a bacterial solution (Clough & Bent, 1998) and a modified procedure for *Agrobacterium* preparation (Logemann *et al*, 2006). T1 plants were selected *in vitro* on the appropriate antibiotic/herbicide and construction integrity were double-checked by genotyping with specific primers (**Appendix Table S4**). Approximately 10 independent T1 plants were transplanted in soil. In the T2 generation, at least two independent lines were selected according the following points: a single insertion of the transgene (Mendelian segregation on selection agent), no obvious abnormal developmental phenotypes, good expression of the transgene for detection by confocal microscopy, with a fluorescence clearly visible and a uniform expression pattern.

### Construct generation for transgenic lines

Plasmids used for plant transformations were generated using the multisite gateway three-fragment strategy (Life Technologies), according the recommendations of the supplier, or by site-directed mutagenesis using PCR amplification of pre-existing plasmids with specific primers (**Appendix Table S4 and S5)**. DNA fragments compatible with entry vectors were generated by PCR amplification of the fragments of interest from genomic DNA with specific primers compatible with Gateway technology or synthesized by Genewiz (**Appendix Table S5**). Amplifications were done with the high-fidelity enzyme Phusion Hot Start II (Thermo Fisher Scientific). PCR products were purified using NucleoSpin® kit (Macherey-Nagel). Plasmids generated, and the corresponding methods are presented in **Appendix Table S5**.

### Pollen observations

*Arabidopsis* fresh pollen was obtained after crushing of young open flowers in liquid solution (water or specific treatment) between blade and coverslip. Fresh maize mature pollen was collected and observed between slide and coverslip in appropriate medium to conserve pollen grain integrity (10% sucrose (w/v), 5 mM CaCl_2_, 1 mM MgSO_4_, 5 mM KCl and 0.01% H_3_BO_4_ (w/v), pH8 with KOH). Nuclear coloration was done using 1 μg/ml DAPI (4’,6-diamidino-2-phénylindole) solution (w/v) (ThermoFisherScientific) diluted in citrate-phosphate buffer (0.1 M C_6_H_8_O_7_, 0.2 M Na_2_HPO_4_ 2H_2_O, pH4 adjustment and addition of 0.1% triton (v/v)). The kinetics of maize pollen development was calibrated on A188 background. Anthers were collected at different developmental stages and classified according to their size and position on the tassel. Anthers were imaged with a binocular loupe to measure their length and then were crushed in DAPI solution between blade and coverslip to evaluate pollen stage development by epifluorescence microscopy (nuclei observation). After calibration, the same experiment was done with *NLD* reporter line to do the temporal analysis of promotor activity.

### Maize sperm cell isolation

Sperm cells were isolated either by osmotic shock or by use of a Percoll gradient, whenever a better purity was needed. For sperm cell isolation by osmotic shock, fresh pollen was placed between slide and coverslip at room temperature in shock medium containing sucrose and low ion concentration (10% sucrose (w/v), 5 μM CaCl_2_, 1 μM MgSO_4_, 5 μM KCl and 0.01% H_3_BO_4_ (w/v), without pH adjustment). For sperm cell isolation by Percoll gradient, 500 mg of fresh pollen was collected in a 15 ml Falcon tube containing 6 ml of 1X BK medium (100X BK: 0.1% H_2_BO_3_ (w/v), 20 mM Ca(NO_3_)_2_, 20 mM MgSO_4_, 10 mM KNO_3_) coupled with 0.52 M mannitol to burst pollen grains. Tubes were vortexed quickly and then gently agitated on a rotary agitator 20 min (4°C). After this step, all the process was done on ice. To remove large debris, the solution was filtered (40 μm and 70 μm) and the filters rinsed with an equivalent volume of 80% Percoll in BK-S15-MOPS medium (1X BK medium, 30% sucrose (w/v), 20 mM MOPS, pH7.5 with NaOH) (w/v). Sperm cell purification was done using 5 ml of filtrate in a new 15 ml tube. Above this volume, 1 ml of 20% Percoll in BK-S15-MOPS medium (w/v) was gently deposited, then 5 ml of BK-Man-MOPS medium (1X BK medium, 0.52 M mannitol, 10 mM MOPS, pH7.5 with NaOH). Tubes were centrifuged 40 min at 1400 rpm (4°C). After this step, two bands containing sperm cells were formed andwere gently removed and individually deposited in new tubes. Six equivalent volumes of BK-Man-MOPS medium were added, and tubes were centrifuged 30 min at 1200 rpm (4°C). Supernatants were mostly removed to leave 500 μl of solution in the tube and the pellets were resuspended in this volume. This suspension containing isolated sperm cells was observed between slide and coverslip.

### Microscopy observations

Maize pollen development analyzes were performed on epifluorecence microscope Imager M2 Axio (Zeiss®), harboring a lamp LED X-Cite® 120LED (Excelitas®). Objectives X20 and X40 were used (Plan-ApoChomat, Zeiss). Confocal imaging of maize pollen marked with both mCitrine and mCherry fluorescent reporters was done on a Leica SP8 up-right confocal microscope, with a water immersion objective (HCX IRAPO L 25x/0.95 W). Fluorophores were excited using Led laser (Leica Microsystems, Wetzlar, Germany) emitting at wavelengths of 514 nm for mCitrine and 552 nm for mCherry. Images were collected at 521–550 nm for mCitrine, 610–650 nm for mCherry. *Arabidopsis* root cells and pollen grain observations for both *Arabidopsis* and maize were done on an inverted Zeiss LSM710 confocal microscope mounted on AxioImager Z2. Samples were observed using a X40 oil objective (Plan-Apochromat 40x/1.4 Oil DIC M27, Zeiss). Dual-colour images were acquired by sequential line switching, allowing the separation of channels by both excitation and emission. Depending on the fluorophores observed, different wavelengths of excitation and band pass filters were used: mCitrine, YFP, VENUS: 514 nm / 520-580 nm and DAPI: 405 nm / 410-480 nm (excitation / band pass). Immunogold labelling imaging was done with a TEM (transmission electron microscopy) Philips CM120 at 120 kV using a CCD camera Gatan Orius 200.

### Immunogold labelling

Pollen was fixed in 2% PFA / 0.2% GA (v/v) in 0.1M Phosphate Buffer (PB) pH 7.4 with 3% sucrose (w/v) under vacuum for 2 h at room temperature and overnight at 4°C. The day after, pollen was rinsed three times with 0.1M PB with 3% sucrose, washed twice in 50 mM glycine (Sigma Aldrich) in 0.1 M PB pH7.4 with 3% sucrose, embedded in 12% gelatin in 0.1M PB with 3% sucrose and placed at 4°C till solid. Small cubes (1 mm^3^ approximately) were cut and then placed in 2.3 M sucrose in 0.1 M PB pH 7.4 ON at 4°C. For cutting, the samples were put on an aluminum pin, snap frozen in liquid nitrogen, and ultrathin sections (100 nm) were cut with a cryo-ultramicrotome (Leica UC S, Leica Microsystems) at −100°C with a glass knife. Sections were picked-up with a mixture of 2% methylcellulose / 2.3 M sucrose (v/v), collected on formvar-carbon-coated copper grids (Electron Microscopy Sciences) and immunostained using standard protocols. Briefly, sections were treated with 50 mM glycine for aldehyde quenching, epitopes were blocked with 0.1 M PBS / 1% Cold Water Fish Skin Gelatin / 0.2% BSA with 3% sucrose (all chemicals from Sigma Aldrich), incubated with the primary antibody anti-GFP (1:250) (XVZ), washed, and then the reaction was revealed using the protein-A-gold (10 nm, 1:70) (Cell Microscopy Core, University of Utrecht).

### PAO and 2-BP drug treatments

Stock solutions of 60 mM PAO (Phenylarsine oxide) in DMSO and 50 mM 2-BP (2-bromopalmitate) in EtOH were ordered from Sigma-Aldrich. Seven-day-old seedling grown *in vitro* vertically (root tip analyses), and fresh open flowers (pollen analyses) were incubated in the dark and under agitation in liquid ½ MS medium complemented with appropriate drug treatment. For roots 60 μM PAO for 30 min or 50 μM 2-BP for 2 h. For open flowers 180 μM PAO for 1 h or150 μM 2-BP for 2 h. For pollen treatment, flowers were wounded with a pincer to facilitate the drug penetration and after treatment they were crushed on the blades to release the pollen grains. The mock conditions correspond to tissue incubation in liquid ½ MS medium supplemented with a volume of DMSO and EtOH, respectively equivalent to the PAO and 2-BP volumes used and for the same time as the actual treatment. Effect of drug treatment was evaluated on roots using confocal image analysis, and on pollen counting directly under the microscope the number of pollen grain from different classes defined according to the subcellular localization of the protein fusion. Three classes were observed: “Endo-PM”: continuous fluorescent signal at the endo-PM, “Spotty”: non-continuous fluorescent signal at the endo-PM, and “Cytosolic”: protein delocalization to the cytosol with low fluorescent signal. For the pollen analyzes five independent treatments were done at the same time.

### Imagining quantification

Image analyzes were performed using the Fiji software. For *Arabidopsis* confocal imaging, different ratios were defined to characterize the subcellular localization of proteins according to the repartition of the fluorescent signal intensity in cells as previously described (Platre *et al,* 2018). Fluorescent intensity of a region of interest (ROI) in pictures was evaluated by the measure of the mean gray value of pixels using the Mean Grey Value function of Fiji software. The ratio of fluorescence PM/cyto was measured on the root cells tips and corresponds to a ratio between the intensity of the fluorescent signal at the level of the plasma membrane (PM) and the intensity of the fluorescent signal at the level of the cytosol (cyto). Per cell, two linear ROI measurements were done (one per cell side) at the level of the plasma membrane and at the level of the cytosol, using the Measure function of software and mean between the both measurements was done. For the transgenic line analyzes, the measurements were carried out on at least five independent lines with the measurement of twenty cells per root tip on at least three different roots per line. For the pharmacological analyzes, the measurements were carried out on three independent experiments, with the measurement of twenty cells per root tips on at least five roots per experiment. The fluorescence ratio endo-PM/PM were measured in mature pollen. It corresponds to a ratio between the intensity of fluorescent signal at the level of the endo-PM and the intensity of the fluorescent signal at the PM level. All pictures for a same experiment were acquired in same microscope setup. For the immunogold labelling quantifications, full sperm cell pictures were reconstituted using the Adobe® Photoshop® software. Gold particles were then quantified by three independent experimenters in a triple-blind set-up by three independent operators (**Appendix Table S3**).

### Statistical analyzes

Statistical analyzes were performed with the RStudio® software. Comparisons of the fluorescence intensity between different samples were performed using the non-parametric Wilcoxon-Mann-Whitney test. Comparison of two different proportions were performed using the chi-square test.

## Acknowledgements

We thank Y. Jaillais and M. Platre for discussions and comments, G. Brunoud, T. Dresselhaus and Y. Jaillais for providing transgenic lines, O. Hamant and Y. Jaillais for critical reading of the manuscript. We are grateful to G. Gendrot, M.-F. Gérentes and J. Laplaige for maize transformation, to J. Berger, P. Bolland and A. Lacroix for maize culture, to I. Desbouchages and H. Leyral for buffer and media preparation, and to J. Guichard for technical assistance. Sample preparation, immuno-electron microscopy experiments as well as transmission electron microscopy observations were performed at the Centre Technologique des Microstructures (CTμ). We acknowledge the contribution of SFR Biosciences (UMS3444/CNRS, US8/Inserm, ENS de Lyon, UCBL) facilities, and in particular C. Lionnet at the LBI-PLATIM-MICROSCOPY for assistance with imaging. This research was supported by the ANR grant (ANR-19-CE20-0012, “Not-Like-Dad”) to T.W., and by the “pack ambition recherche” from the Région Auvergne-Rhone-Alpes (“DH-INNOV”) to T.W. N.M.A.J. and L.M.G. were supported by CIFRE PhD fellowships from ANRT funding agency (grant # 2019/0771 and 2015/0777, respectively).

## Author contributions

LMG and TW conceived and designed the experiments. LMG, NMAJ, VL and TW conducted experiments. VL did immuno-EM and TEM imaging. LMG prepared tables and figures. LMG and TW led the writing of the manuscript and VL, PMR, JPM contributed to the critical reading of the manuscript. PMR, JPM and TW were involved in project management. TW obtained funding. TW initiated and coordinated the project.

## Tables and their legends

NA

## Expanded View Figure legends

**Expanded View 1.**
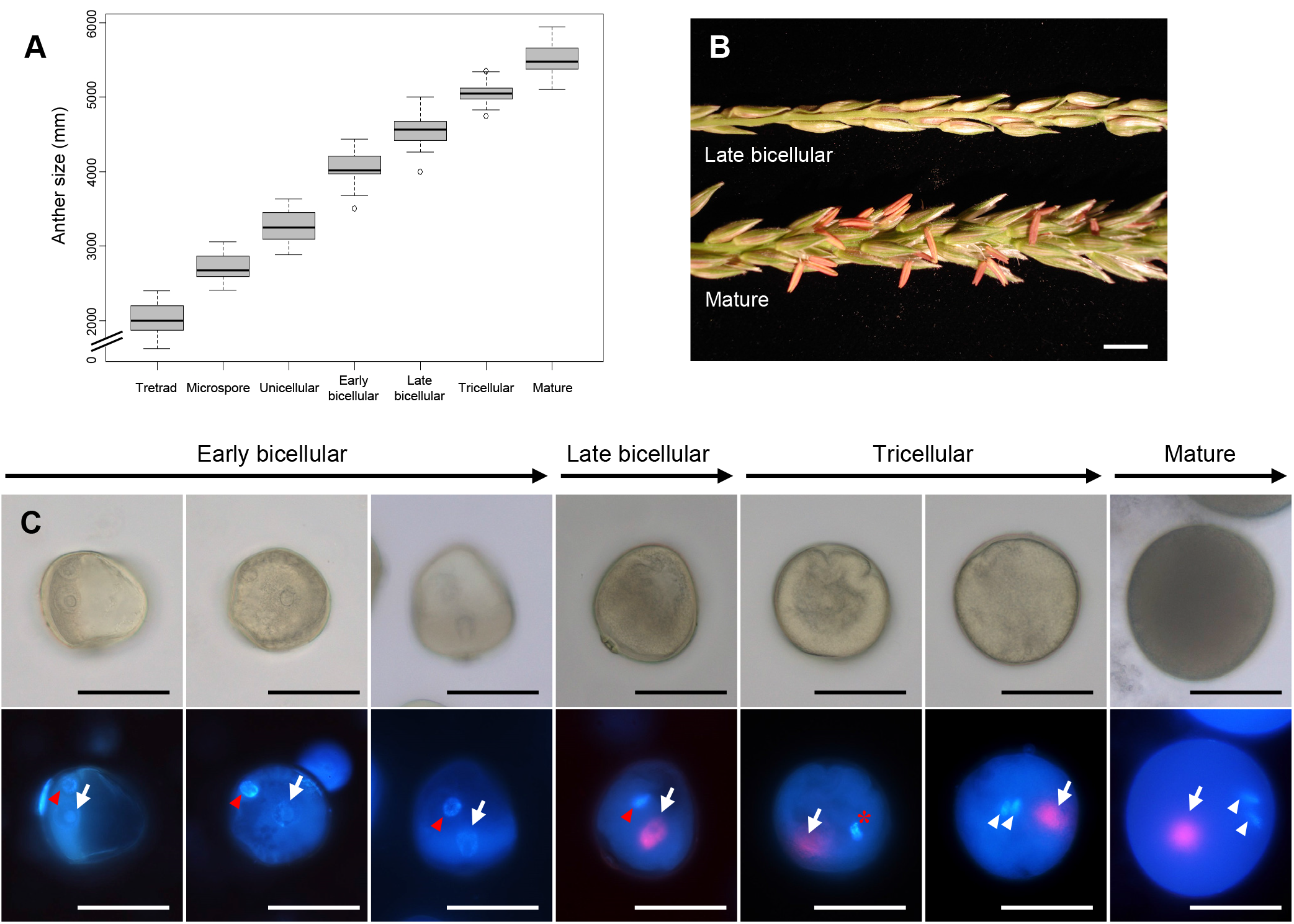
*NLD* promoter activity in the vegetative cell starts at the late bicellular stage, just before formation of the two sperm cells. **A**: Calibration of anther size according to pollen development stages in A188 background. 36 anthers were measured for each stage. **B**: Tassel branches with male flowers at late bicellular stage (top picture) when *NLD* promoter activity is starting, and mature pollen stage (bottom picture). **C, D**: Temporal analysis of *NLD* promoter activity (*pNLD::H2B:mCherry*) at different stages of pollen development, between early bicellular and mature pollen stages. *NLD* promoter activity (red signal) starts at late bicellular stage, probably just prior to the second pollen mitosis. White arrows point at the vegetative nucleus, red arrow heads at the generative cell nucleus, and white arrow heads at the sperm cell nuclei. The red asterisk indicates a second pollen mitosis in progress. Determination of pollen stages by epifluorescence imaging of pollen after DAPI (4’6-diamidino-2-phenylindole) staining. Scale bars: B = 500 μm, C, D = 50 μm.

**Expanded View 2:**
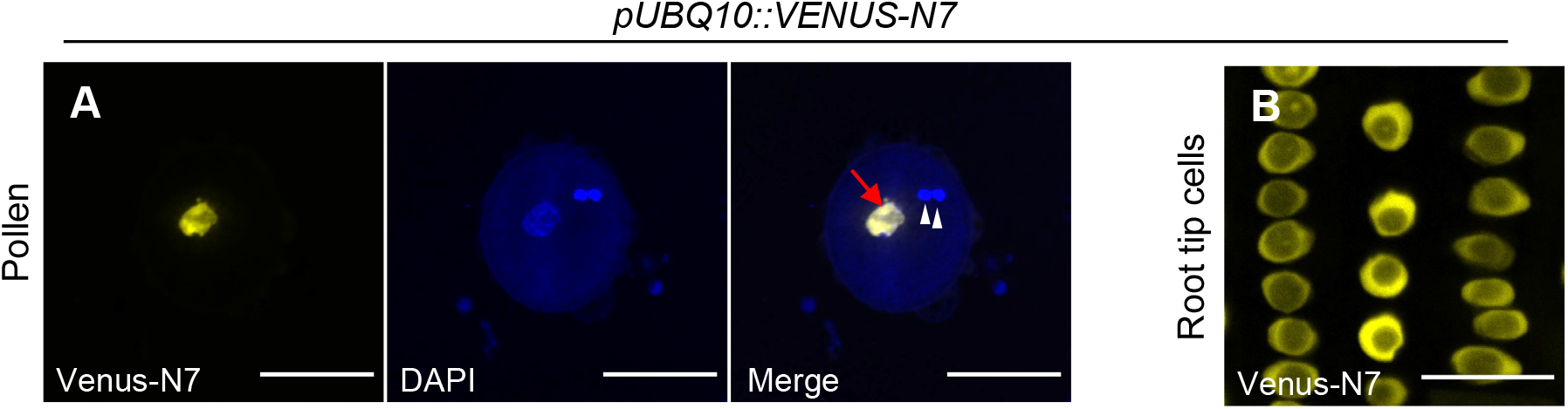
Validation of the *pUBQ10* promoter expression pattern in *Arabidopsis* pollen. **A, B.** Confocal imaging of mature pollen (A) and root apex (B) of the *pUB10::Venus-N7* line (yellow signal). DAPI (4’6-diamidino-2-phenylindole) stains nuclei (blue signal), white arrow heads indicate the sperm cells nuclei, and red arrow points at the vegetative nucleus.

**Expanded View 3:**
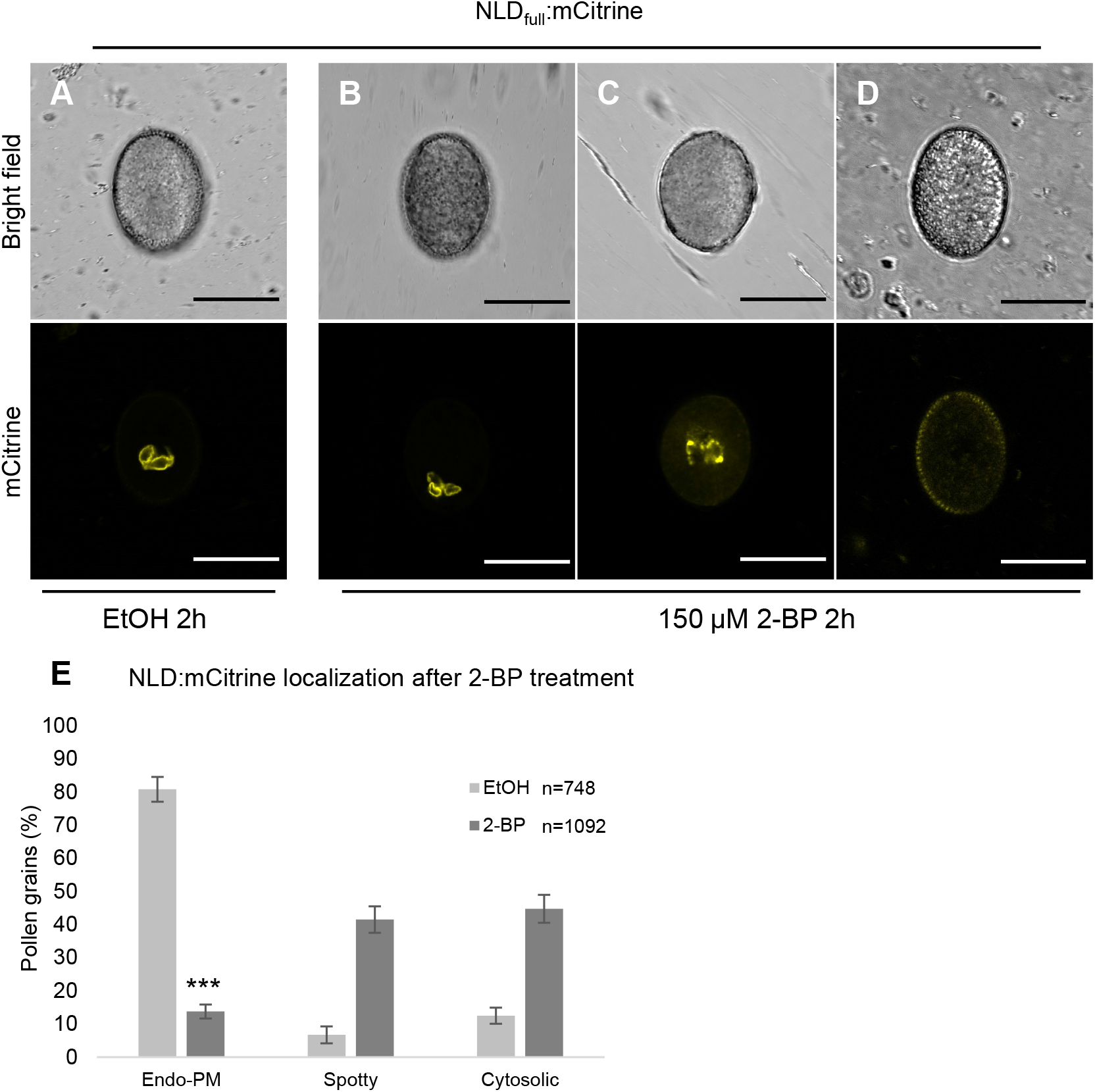
Pharmacological approach showing the involvement of S-palmitoylation in NLD attachment to the endo-PM in *Arabidopsis* pollen. **A - D**. Confocal imaging of mature pollen expressing NLD_full_:mCitrine after 2 h treatment **A**: with mock (EtOH) or with (**B-D**) 150 μM 2-BP (2-bromopalmitate, an inhibitor of palmitoylation). After mock treatment, NLD_full_:mCitrine mainly localizes at the endo-PM (A), whereas 2-BP treatment leads to three different types of localization: Endo-PM (B), non-continuous (“spotty”) fluorescence signal at endo-PM (C) and protein delocalization to the cytosol with low fluorescent signal (D). Scale bars = 20 μm. **E** Quantification of drug treatment effect on NLD subcellular localization by counting the three different classes of localization patterns after 2-BP treatment. *** denotes significant difference (p-value < 0,001; Chi-square test, n = number of counted mature pollens).

**Expanded View 4.**
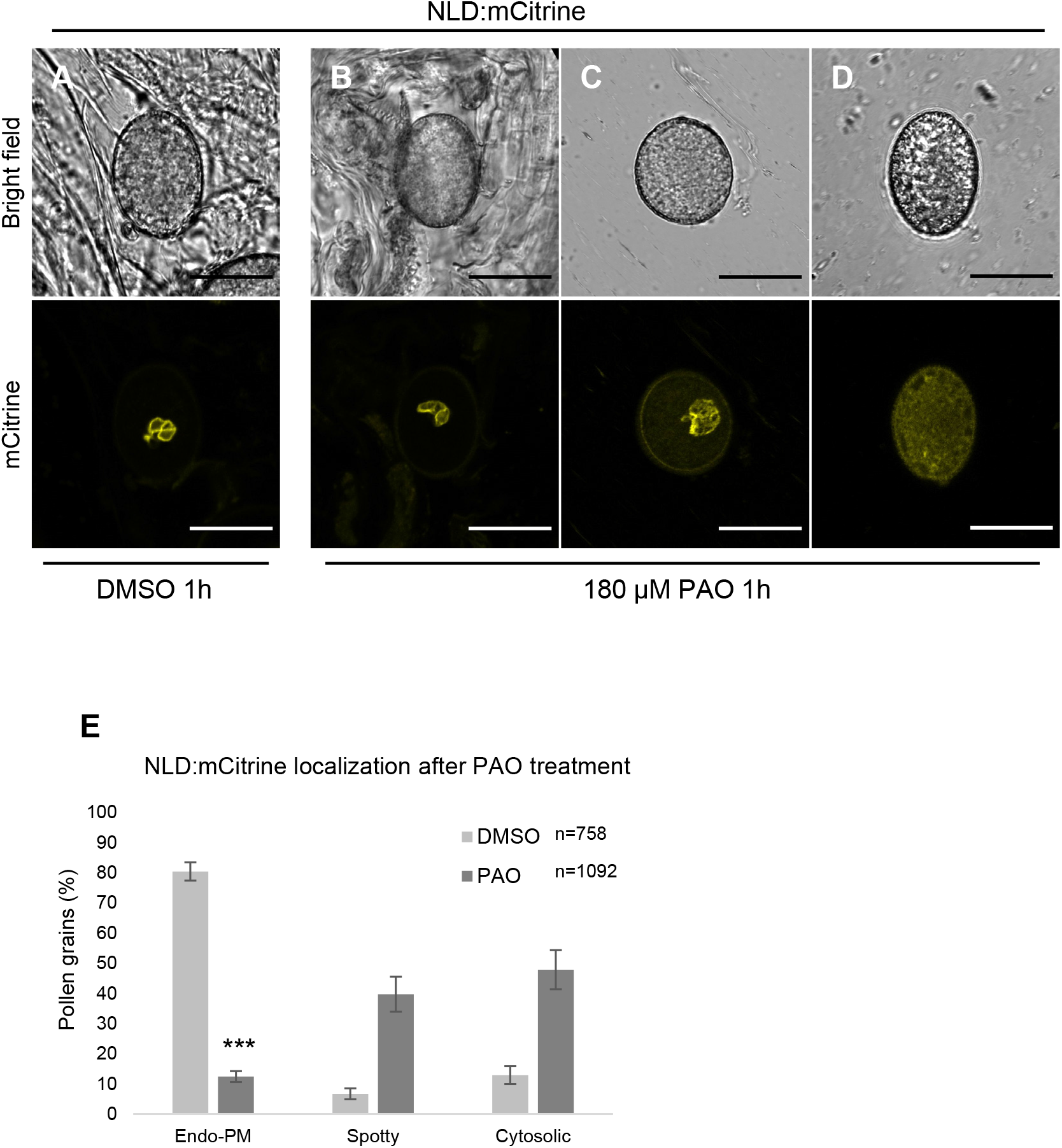
Pharmacological approach showing the involvement of electrostatic interactions between NLD and endo-PM in pollen. **A - D** Confocal imaging of mature pollen from lines expressing NLD_full_:mCitrine after 1 h treatment with mock (DMSO) or with 180 μM PAO (Phenylarsine oxide), an inhibitor of PI4-Kinases. After DMSO treatment, NLD_full_:mCitrine localized to the endo-PM (A, E), whereas PAO treatment led to three different types of localization: Endo-PM (B), non-continuous (“spotty”) fluorescence signal at endo-PM (C) and protein delocalization to the cytosol with low fluorescent signal (D). **E**: Quantification of drug treatment effect on NLD subcellular localization by counting the three different classes of localization patterns after PAO treatment. *** denotes significant difference (p-value < 0,001; Chi-square test, n = number of counted mature pollens).

## Appendix Tables

**Appendix Table S1**: Description of maize and Arabidopsis transgenic lines used in this study

**Appendix Table S2**: Complementation of haploid induction phenotype by *pNLD::NLD:mCitirne*

**Appendix Table S3**: Quantification of gold particles indicating NLD:mCitrine localization.

Counting of gold particles observed after immunogold labeling of *pNLD:NLD:mCitrine* and non-transgenic pollen grains, by three different operators. Vegetative cell: particles found within the part of the vegetative cell that is visible on the reconstituted picture. Endo-PM: particles found in the Endo-PM. Interspace: particles found within the space between sperm cell PM and endo-PM; Sperm cell: particles found in sperm cells (cytoplasm, nucleus and sperm cell-PM). NA: particles that could not be assigned to a particular compartment.

**Appendix Table S4**: List of primers used in this study

**Appendix Table S5**: Detailed of plasmids and clonning methods used in this study

